# Nicotinic signaling is required for motor learning but not rehabilitation after spinal cord injury

**DOI:** 10.1101/2022.01.13.476174

**Authors:** Yue Li, Edmund R Hollis

## Abstract

Currently, therapeutic intervention for spinal cord injury is limited. Many approaches rely on strengthening the remaining substrate and driving recovery through rehabilitative training. As compared to learning novel compensatory strategies, rehabilitation focuses on restoring movements lost to injury. Whether rehabilitation of previously learned movements after spinal cord injury requires the molecular mechanisms of motor learning, or if it engages previously trained motor circuits without requiring novel learning. Our findings implicate the latter mechanism, as we demonstrate that nicotinic acetylcholine signaling is required for motor learning but is dispensable for the recovery of previously trained motor behavior after cervical spinal cord injury.

## Introduction

Spinal cord injury results in the lasting impairment of the motor and sensory functions that underlie movement. The majority of clinical cases of spinal cord injury are incomplete, allowing for a limited capacity for spontaneous or rehabilitation-mediated functional recovery. This capacity for partial recovery may be through reinforcement of spared sensory and motor axons, compensatory activities of indirect pathways, or reorganization of supraspinal command centers (Hollis et al., 2016; Li and Hollis, 2017). Critical to restoring supraspinal command of voluntary movement is the corticospinal tract. Following stroke, the extent of spared corticospinal tract is proportional to the amount of spontaneous recovery that individuals experience (Stinear et al., 2007). Rehabilitation and epidural electrical stimulation approaches to restore voluntary movements after spinal cord injury likely act in part upon remaining corticospinal axons (Wagner et al., 2018). These spared circuits can be activated by cortical stimulation even in motor complete, chronically injured individuals (Edwards et al., 2013).

Both rodent and non-human primate models have been used to demonstrate the innate plasticity of corticospinal axons after injury (Rosenzweig et al., 2010; Mosberger et al., 2017). Within the injured spinal cord, corticospinal axons sprout locally and form novel axon collaterals, many of which are pruned back over time (Bareyre et al., 2004). The removal of intrinsic brakes on axon regeneration can enhance corticospinal axon plasticity and connectivity; however, the contribution of such connections to behavioral recovery is not always apparent (Liu et al., 2010; Hollis et al., 2016; Jayaprakash et al., 2016). Rehabilitative training drives recovery of previously trained, corticospinal-dependent, single pellet reach behavior in animal models of corticospinal tract injury (Wahl et al., 2014; Hollis II et al., 2016). Clinically, training after injury can be used to either rehabilitate movements lost to injury, or to train compensatory strategies to improve mobility and independence (Behrman and Harkema, 2007).

Motor learning requires contributions from various brain areas, including motor cortex, cerebellum, striatum, and brainstem. Basal forebrain cholinergic neurons release acetylcholine in distinct targets and modulate a diverse array of functions, such as motor control, attention, cognition, and perception coding (Zaborszky et al., 2018; Boskovic et al., 2019). The cerebral cortex receives cholinergic input from nucleus basalis of Meynert (NBM). Primary motor cortex (M1) depends upon these basal forebrain cholinergic neurons for the maturation of cortical motor representations, or motor maps (Ramanathan et al., 2015). Ablation of NBM cholinergic neurons in rats attenuates skill acquisition on the single pellet reach task as well as the corresponding expansion of cortical forelimb motor representations and dendritic spine remodeling of corticospinal neurons that control the distal forelimb (Conner et al., 2003; Wang et al., 2016). Following cortical injury, rehabilitative training of skilled forelimb movements results in a reconstitution of affected movement representations within adjacent, ectopic cortical areas (Castro-Alamancos et al., 1992; Castro-Alamancos and Borrel, 1995). As with motor learning, cholinergic input is critical for motor map reorganization and functional recovery after cortical injury (Friel et al., 2000; Conner et al., 2005). Previously, we observed similar cortical reorganization alongside functional recovery after spinal cord injury (Hollis et al., 2016). This recovery was dependent on rehabilitative training. Unlike following stroke, cortical structures remain intact after spinal cord injury. It remains unknown whether the molecular and cellular mechanisms supporting motor learning are required for rehabilitation-mediated behavioral recovery, or if rehabilitation relies on reactivation of previously trained motor circuits.

Nicotinic acetylcholine receptors expressed in the central nervous system are important for synaptic excitation, attention, and cognition (Dani, 2001). In primary visual cortex, cholinergic innervation is required for experience-dependent plasticity during the critical period (Bear and Singer, 1986) and closure of the critical period is associated with reduced nicotinic signaling (Morishita et al., 2010). Here we describe the effects of nicotinic cholinergic signaling on skilled motor task acquisition and rehabilitation after spinal cord injury.

### Materials and Methods Animals

All animal experiments and procedures were approved by the Weill Cornell Medicine Institutional Animal Care and Use Committee. All mice were housed on a 12-hour light/dark cycle from 6am to 6pm at 25°C with free access to food and water. Male and female C57BL/6 animals (8-12 weeks old) were purchased from Jackson Laboratory. For forelimb reach task, animals were food restricted to 80–90% of their free-feeding bodyweight.

### Drug administration

Mice were injected with Mecamylamine (MEC, 5 mg/kg, IP; Tocris, 2843) and Methyllycaconitine citrate salt (MLA, 5 mg/kg, IP; Sigma-Aldrich, M168) or vehicle (normal saline) 30 min before behavioral testing.

### Recessed single pellet reach task

We employed a recessed single pellet reach task as described (Li and Hollis, 2021). Animals were calorie restricted to 80–90% of their free-feeding bodyweight before training. An acrylic behavior box with three slots (7 mm wide) on the left, middle, and right sides of the front wall was used to train the mice. A recessed hole (3 mm wide, 2 mm deep) at 12 mm from the inside wall of the box was used to hold a 20-mg flavored food pellet (Bioserv, F05301). The dominant forelimb was identified during a single test session. Once the dominant forelimb was determined, it was trained over a total of 14 daily sessions consisting of 25 trials each. A trial was counted as a success if the mouse grasped, retrieved, and ate the food pellet. Only trials with pellet contact were counted. The intensive rehabilitative training was performed one week after spinal cord injury. Animals were trained 25 trials daily and 15 sessions in total. The success rate was defined as the percentage of trials with successful retrieval and eating.

### Rotarod test

Mice were habituated on the rotarod (Med Associates ENV-577M) at a speed of 4 rpm for 60 s before testing. For each trial, the rotarod accelerated from 4 to 40 rpm over 300 s, then remained at 40 rpm for an additional 300 s, as necessary. Animals were tested 30 min after intraperitoneal injection of nicotinic receptor inhibitors MEC and MLA or vehicle, one trial was performed per day. The latency to fall after the onset of acceleration during each trial was recorded for each mouse. Individual trials were stopped, and the duration was recorded, if mice could not run with consecutive rotations or failed to stay on the rotarod. If animals successfully completed 600 s on the rotarod, the latency was recorded as 600 s.

### Open field test

To test animal activity, mice were placed in a chamber (length × width × height: 30 × 22.5 × 25 cm) and allowed to explore for 5 min. Behavior was recorded from the top at 48 fps (GoPro, HERO3) and analyzed by MATLAB software (Autotyping15.04) (Patel et al., 2014).

### C5 dorsal column lesion

Mice were anesthetized with 4% isoflurane, maintained during surgery with 1.5–3% isoflurane, and had body temperature maintained at 37°C using a SomnoSuite small animal anesthesia system (Kent Scientific). Subcutaneous injection of buprenorphine (0.1mg/kg) was given immediately following anesthesia. Spinal level C5 was exposed by laminectomy and the dorsal columns were lesioned at a depth of 1 mm with Vannas spring scissors, as previously performed (Hollis II et al., 2016). The dorsal musculature was sutured with 6-0 suture and the skin was closed with wound clips. Mice were allowed to recover from surgery for one week before rehabilitative training.

### Histology

To confirm spinal cord injury, after finishing all behavioral training, animals were anesthetized with ketamine/xylazine cocktail (150mg/kg; 15mg/kg) and transcardially perfused with ice-cold PBS followed by 4% paraformaldehyde (PFA). Spinal columns were postfixed in 4% PFA overnight at 4°C, cryoprotected by immersion in 30% sucrose in 0.1 M PBS for 2 d. Samples were then sagittally cryosectioned at 20 μm using a Leica cryostat and directly mounted on Superfrost Plus slides (Fisher Scientific). Sections were blocked with 10% donkey serum for 1 h, and incubated with rabbit anti-PKCγ (1:100, sc-211, Santa Cruz Biotechnology; RRID:AB_632234) and mouse anti-GFAP (1:750, ab10065, Abcam; RRID:AB_296804) antibodies for 2 d at 4°C. Sections were then washed three times with PBS and incubated with fluorescently conjugated donkey anti-rabbit/mouse secondary antibody (1:200, Jackson ImmunoResearch) for 1.5 h at room temperature. Images were acquired on a Leica SP8 confocal microscope with 10X objective.

### Statistical analysis

Skilled pellet reach and rotarod tests were analyzed using two-way repeated measures ANOVA with post hoc Sidak’s comparison test using GraphPad Prism 9.0. The differences between two groups were compared by two-tailed t-tests. For all figures, *P < 0.05, **P < 0.01, ***P < 0.001.

## Results

### Systemic inhibition of nicotinic receptors impairs motor learning

We used a pharmacological approach to study the contribution of nicotinic receptors to motor learning in adult mice. Mecamylamine (MEC, 5 mg/kg body mass) and methyllycaconitine (MLA, 5 mg/kg) were injected intraperitoneally (IP) 30 min prior to behavioral training. Control mice were injected with normal saline. Single pellet reach is a skilled behavior used to measure dexterity of a single forelimb (Whishaw and Pellis, 1990). We employed a modified recessed single pellet reach task in which the food pellet is retrieved from a concave depression; we previously found that this modification allows for a consistent learning curve in C57BL/6J mice (Li and Hollis, 2021). We found that systemic blockade of nicotinic receptors significantly attenuated skilled motor learning on the single pellet reach task (Figure 1A,B). MEC and MLA-treated mice showed smaller improvements over the course of training than controls. Washout of MEC and MLA over 5 days enabled the mice to learn the task with the same proficiency as controls (Figure 1A,B).

**Figure 1.**
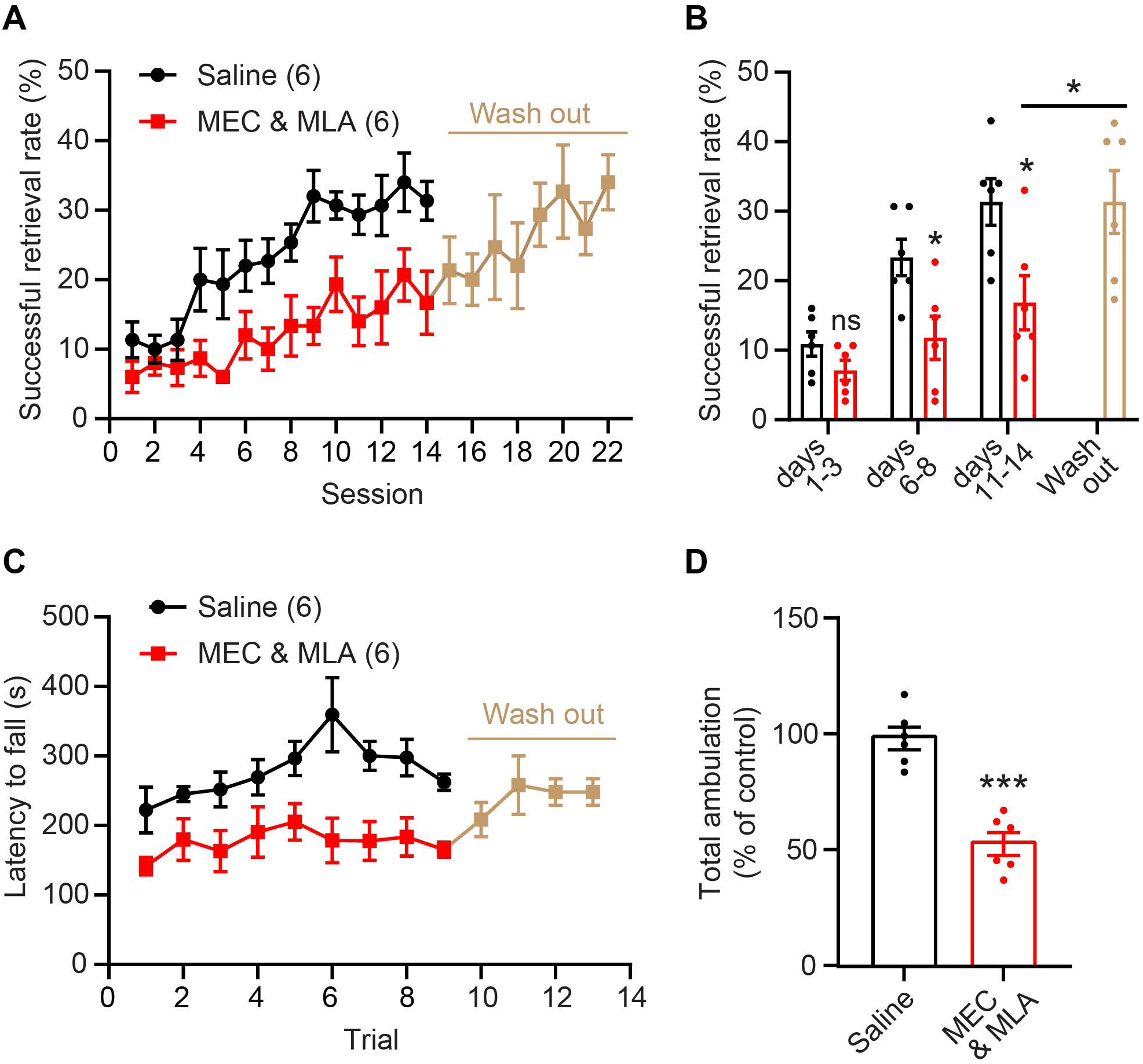
Inhibition of nicotinic acetylcholine signaling impairs motor learning. (**A**) IP injection of MEC and MLA 30 minutes prior to training significantly attenuated learning on the recessed single pellet reach task (*P* = 0.0086, repeated measures ANOVA, F(1,10) = 10.6). After 14 days of training, MEC and MLA treatment was stopped. Five days later, mice were trained for an additional 8 days without treatment. (**B**) Mean value analysis. Day 1-3 is the initial phase of motor learning, day 6-8 is the rising phase, and day 11-14 is the plateau phase. There was no effect of MEC and MLA treatment during the initial phase, but the mean values during the rising and plateau phases were significantly lower in drug-injected animals than that in controls (\**P* < 0.05, unpaired t-test). Following wash out of MEC and MLA, the mice were able to learn the task (mean success rate over days 19-22 vs days 11-14, \**P* < 0.05, paired t-test). (**C**) Administration of MEC and MLA impaired rotarod training performance (*P* = 0.0067, repeated measures ANOVA, F(1,10) = 11.62). Following wash out, mice were able to learn the rotarod task. (**D**) Total walking distance was shorter in mice injected with MEC and MLA (n = 6/group, \*\*\**P* < 0.001, unpaired t-test).

The effects of systematic inhibition of nicotinic acetylcholine signaling were not specific for skilled forelimb motor learning. MEC and MLA administration also impaired coordinated motor learning on the accelerating rotarod task. Mice trained over nine trials (4 to 40 rpm, constant acceleration over 5 min) exhibited worse performance following IP injection with MEC and MLA compared to control mice. MEC and MLA inhibition of nicotinic signaling resulted in significantly reduced latency to fall (Figure 1C). As with single pellet reach, washout of MEC and MLA over 5 days enabled the mice to learn the rotarod with the same proficiency as controls (Figure 1C). Additionally, MEC and MLA significantly reduced total walking distance in an open field test (Figure 1D).

### Nicotinic signaling is not required for functional recovery following spinal cord injury

To test whether nicotinic signaling is also required for functional recovery, we randomly reassigned animals after motor learning, performed a cervical spinal cord injury, and tested the effects of MEC and MLA on intensive rehabilitative training on the single pellet reach task. We performed a dorsal column lesion at cervical spinal cord segment 5 (C5) to transect the corticospinal and ascending dorsal column tracts, leaving most gray matter, lateral white matter, and ventral spinal cord intact (Figure 2A). One week after C5 dorsal column lesion, intensive rehabilitative training was carried out in which animals were tested daily on the recessed pellet reach task. Spinal cord injury significantly impaired task success (Figure 2B). Intensive rehabilitative training promoted recovery to levels similar to pre-injury, regardless of treatment group; MEC and MLA delivery had no effect on the recovery of function (Figure 2B).

**Figure 2.**
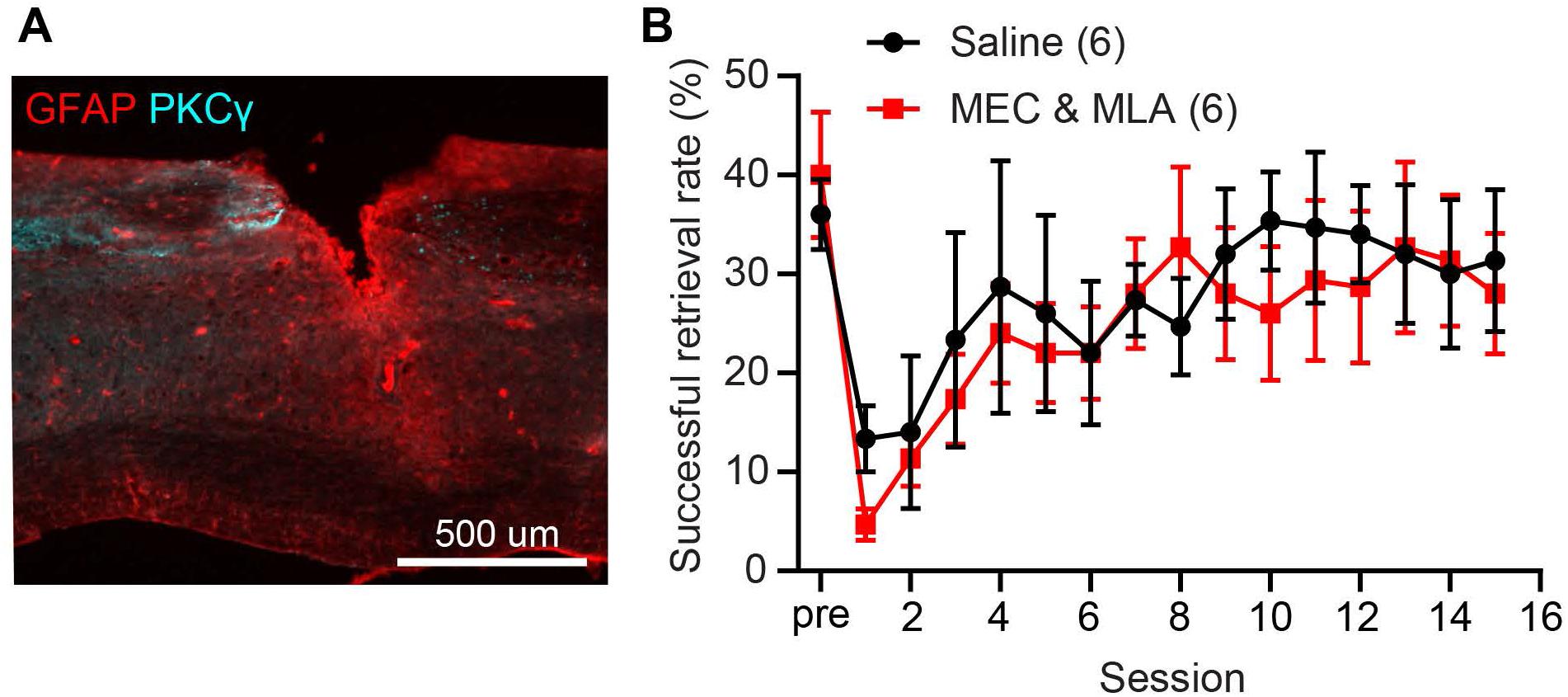
Inhibition of nicotinic receptors do not prevent functional recovery after spinal cord injury. (**A**) Representative image of injured cervical spinal cord with PKCγ-labeled corticospinal axons and GFAP-labeled astrocytes. (**B**) Animals injected with MEC and MLA show similar functional recovery on the single pellet reach behavior to vehicle-treated control animals (*P* = 0.76, repeated measures ANOVA, F(1,10) = 0.1026).

## Discussion

In this study, we used pharmacological tools to demonstrate that nicotinic acetylcholine signaling is required for the acquisition of motor skills but not the rehabilitation-mediated recovery of the previously trained skills after spinal cord injury. Recently we have found that mice, unlike rats, do not require basal forebrain cholinergic input to cortex for the acquisition of skilled motor learning on the forelimb reach task (Li and Hollis, 2021). This leaves open the question of the locus of nicotinic activity during motor learning. In our previous study, we targeted both NBM cholinergic neurons directly, through targeted toxin, genetic, and optogenetic means, as well as the projections of NBM cholinergic neurons to motor centers in medial prefrontal cortex and primary motor cortex, leaving cholinergic innervation of striatum, brainstem, cerebellum, spinal cord, and periphery intact. Nicotinic signaling is likely active in one of these other motor loci during motor learning.

During rehabilitation from spinal cord injury, the extent to which rehabilitation either relies on the execution of previously encoded motor programs or else leverages the cellular and molecular mechanisms of motor learning is not known. Motor cortex plasticity occurs alongside the acquisition of skilled motor learning and we previously found that rehabilitation on the single pellet reach task after spinal cord injury shapes both behavioral recovery and cortical plasticity (Hollis II et al., 2016). During motor learning, the role of motor cortex diminishes commensurate with the development of task proficiency. Inactivation of primary motor cortex early in training of a forelimb lever press task impaired performance; however, cortical silencing after an extensive training period had little effect on task success or movement kinetics (Hwang et al., 2019). In fact, the execution of a similar trained temporally precise lever press task is essentially unperturbed by the bilateral aspiration of the entire motor cortex (Kawai et al., 2015). The declining role of motor cortex in execution of learned behavior is reflected in the absence of a role for cholinergic signaling following training. We previously found that ablation of cholinergic innervation of motor cortex after coordinated motor learning of rotarod behavior had no effect on task execution (Li and Hollis, 2021), similar to the absence of effects on single pellet reach task success in rats when cholinergic neurons were ablated after training (Conner et al., 2003). Thus, when animals become proficient in a motor skill, motor cortex disengages from the behavior and subcortical structures (such as basal ganglia, red nucleus, brain stem, and cerebellum) are sufficient for maintenance of previous learned motor skills (Hikosaka et al., 2002).

The spinal cord receives multiple motor inputs and these supraspinal circuits are likely to control different aspects of movement execution. Our dorsal column spinal cord injury was limited to transection of the main body of the descending corticospinal tract and the ascending dorsal column-medial lemniscal sensory circuit, leaving other supraspinal pathways intact, including rubrospinal, reticulospinal, and the lateral, minor corticospinal tracts. It may be that the remaining supraspinal motor circuits retain the motor patterns encoded through training needed to compensate for the loss of dorsal column circuitry, or that nicotinic signaling plays no role in the shaping of these alternate motor pathways. Others have implicated a role for spared ventral corticospinal axons in rats and dorsolateral corticospinal axons in mice in the restoration of corticospinal-dependent behaviors, with limited effect on cortical motor representations (Weidner et al., 2001; Hilton et al., 2016). While we found no role for nicotinic signaling during intensive rehabilitative training to restore a previously trained, stereotyped movement, nicotinic signaling is likely to play a role in the learning of novel, compensatory movement strategies employed by individuals to regain independence after spinal cord injury.

## Author contributions

YL and EH designed the study and wrote the manuscript. YL performed the experiments and analyzed data.

## Acknowledgements

We acknowledge the Burke Neurological Institute Structural and Functional Imaging Core.

## Data availability

The complete dataset is available at doi.gin.g-node.org/10.12751/g-node.atjjnn/

